# Response diversity increases functional stability but decreases diversity and compositional stability of grassland communities

**DOI:** 10.1101/2024.08.08.607162

**Authors:** Vincent Zieschank, Robert R. Junker

**Affiliations:** Evolutionary Ecology of Plants, Department of Biology, Philipps-University Marburg, 35043 Marburg, Germany

**Keywords:** grass sods, land-use intensity, performance-environment relationships, quantification of asymmetric dependence *qad*, response dissimilarity, ecosystem stability

## Abstract

The insurance hypothesis of biodiversity assumes that ecosystem stability rises with increasing biodiversity because functionally redundant species respond differently to environmental changes, allowing some species to compensate for the loss of others. We tested this hypothesis by combining extensive field data and a common garden experiment where sods originating from different regions were subjected to land-use treatments. Based on plant species-specific performance-environment relationships with abundance as performance proxy and land-use intensity as environmental variable, we calculated response dissimilarity of species-pairs. The resulting dissimilarity matrix was used to calculate response diversity (functional dispersion) of grass sods before and after land-use treatments. Our results showed that high land-use intensity decreased response diversity of plant communities both in the field as well as in the common garden. Response diversity in grass sods increased functional stability but decreased stability in terms of species diversity and composition as communities with high response diversity lost species without replacement in response to experimental land-use change, while those with low response diversity showed species turnover. We conclude that response diversity is an important component of biodiversity and discuss future research directions to refine and generalize the concept of response diversity and its role in ecosystem stability.

## Introduction

Community responses to anthropogenic perturbations define ecosystem stability and thus long-term functionality. It is well established that biodiversity is key for communities’ resistance and resilience against global change components and thus the maintenance of important ecosystem functions (Hautier et al., 2015; Tilman et al., 2006; Wang et al., 2019). The intensification of land-use is among the most severe threats for grasslands (Díaz et al., 2019; Foley et al., 2005; Newbold et al., 2015), which poses stress on these plant communities that negatively respond to mowing and/or fertilization (Allan et al., 2015; Socher et al., 2013; Wesche et al., 2012). As land-use intensifies, some plant species thrive (winners) while others suffer (losers) (Boulangeat et al., 2012; Busch et al., 2019; Völler et al., 2017). This leads to an increasing homogenization of plant communities and destabilizes communities as well as ecological networks and associated functions (Felipe-Lucia et al., 2020; Gossner et al., 2016; Manning et al., 2015). The insurance hypothesis of biodiversity – as one potential mechanism underlying the link between biodiversity, stability, and function (Mori et al., 2013; Yachi & Loreau, 1999) – proposes that biodiverse communities may functionally compensate the loss of some species and thus prevent ecosystem degradation (Bazzichetto et al., 2024; Blüthgen et al., 2016; Dıíaz & Cabido, 2001; Lisner et al., 2023). A prerequisite for this compensation are heterogenous responses to environmental changes of functionally redundant species within a community (Mori et al., 2013; Nash et al., 2016). Accordingly, ‘response diversity’, defined as interspecific differences in responses to abiotic factors, has been identified as an important but understudied component of biodiversity that facilitates ecosystem stability (Cariveau et al., 2013; Ross & Sasaki, 2024; Sasaki et al., 2019). While the potential of response diversity in explaining the relation between biodiversity and ecosystem stability is widely accepted and debated in recent literature, the observational, experimental, and statistical methods required to quantify response diversity did so far not converge to best practices (Ross et al., 2023; Ross & Sasaki, 2024). For instance, the use of response trait diversity as proxy for response diversity (Aquilué et al., 2020; Craven et al., 2016) has been criticized for the difficulty in discriminating response from effect traits responsible for uncertainties in the interpretation of results (Mori et al., 2013; Ross et al., 2023). More promising may be ‘performance-environmental relationships’ (Fig. 1A, Ross et al., 2023) that directly reflect species responses to environmental changes. Based on performance proxies of several species along the same environmental gradient, the interaction term *species x environment* has frequently been considered as indicative for response diversity (Stavert et al., 2017; Winfree & Kremen, 2009). Alternatively, the comparison of performance curves of species pairs is a promising approach that reveals more detailed information on species’ responses and their differences to the environment (Fig. 1B, Ross et al., 2023).

**Fig. 1.**
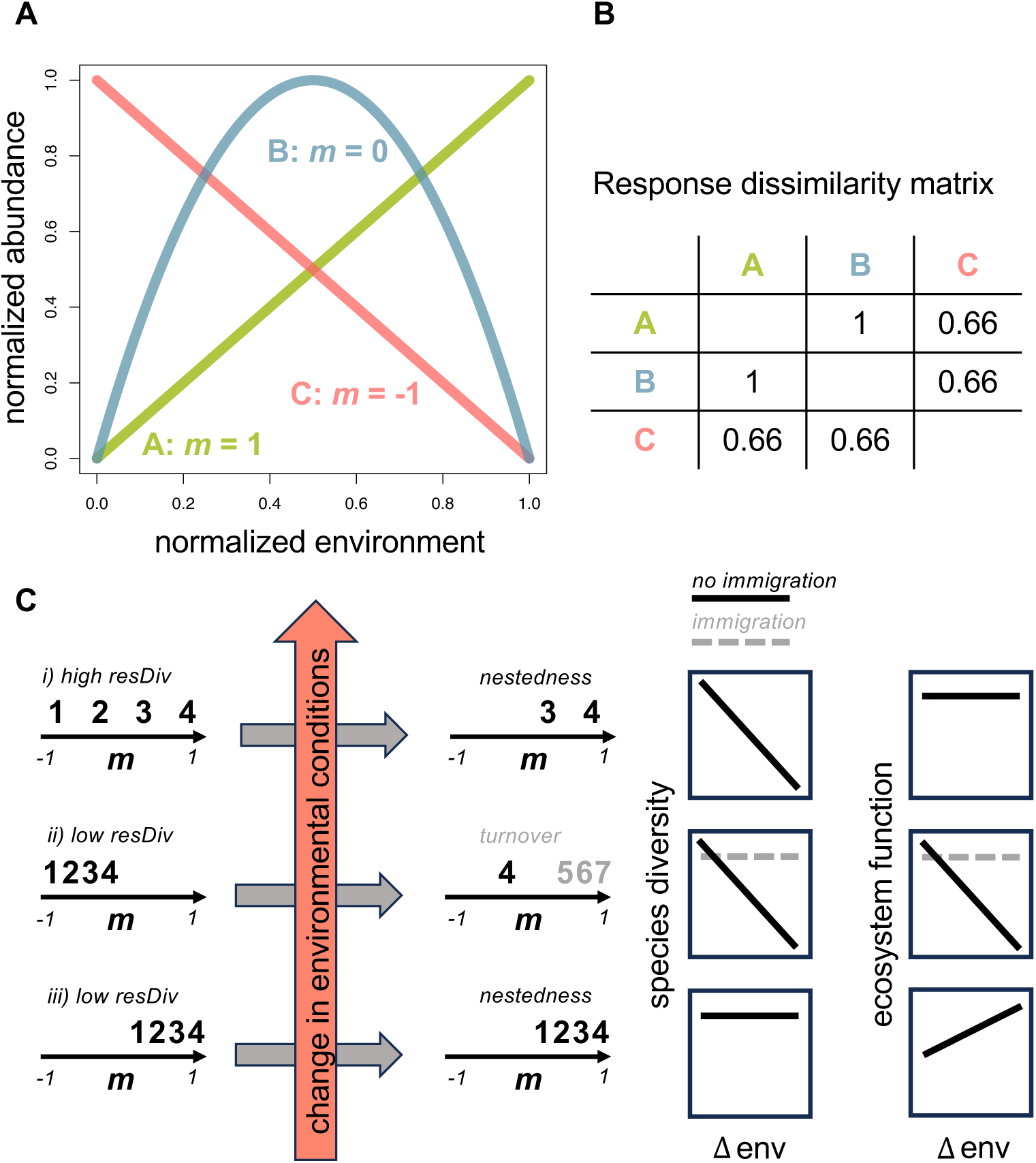
Expected species and community responses to changing environments in relation to response dissimilarity and diversity. Changing environments may include global change components such as warmer temperatures or increasing land use, or natural gradients such as elevation, successions, or latitude. **A)** Abundance-environment relationship of three species A (monotonic positive response to increasing environment values), B (non-monotonic response to environment values), and C (monotonic negative response to increasing environment values). The index of monotonicity *m* defined in Hanusch et al. (2023) measures the deviation from a perfect monotonic positive and negative response to the environment, i.e. *m* = 1 indicates a perfect monotonic positive association to the environment; *m* = -1 a perfect monotonic negative association; and *m* ≈ 0 a non-monotonic association. **B)** Response dissimilarity ∈ [0, 1] quantifies pairwise distances between species-specific abundance-environment relationships described by two empirical checkerboard copulas that are two-dimensional distribution functions that contain full information about associations (Griessenberger et al., 2022; Junker et al., 2021). Responses of A and B to the environment represent the two extremes (perfect monotonic positive and negative responses, respectively) and thus obtain the value 1. Response dissimilarity matrix can be used to quantify response diversity RD of communities analogous to distance-based functional diversity measures (Laliberté & Legendre, 2010; Villéger et al., 2008). **C)** Expected community responses to changing environments in terms of species composition and diversity as well as ecosystem function dependent on initial response diversity RD. Species (1, 2, …, 7) in each community (i, ii, iii) are characterized by the index of monotonicity *m* informing about their responses to environmental changes (see A). Thus, communities with species covering the whole response gradient from negative (*m* = -1) over non-monotonic (*m* = 0) to positive responses (*m* = 1) feature a high response diversity (i), whereas communities with species covering a part of the response gradient only (ii and iii) feature a low response diversity. After positive changes in the environment (indicated by red arrow and ‘Δ env’, i.e. after climate warming or increased land use intensity), we expect the following consequences for communities: i) Community with high response diversity and species that positively respond to positive changes in the environment (species 3 and 4; *m* > 0) are well adapted to the new environment and will dominate the communities, which will lead to a reduced species diversity but a constant ecosystem function as predicted by insurance hypothesis (Yachi & Loreau, 1999). The new community will have a reduced response diversity and species present in the new community will be nested in the species of the previous community. ii) Communities with low response diversity and species with on average negative responses to positive environmental changes (*m* < 0) will suffer from reduced species and response diversity as well as decreasing ecosystem function (black solid lines). Immigrating species (5, 6, and 7, adapted to new environmental conditions) may compensate losses in species diversity as well functionality (grey dashed lines). In these cases, we would expect a turnover in species composition. iii) Communities with low response diversity and species with on average positive responses to positive environmental changes (*m* > 0) will retain species and (low) response diversity and may even increase their functionality. New species composition will be nested in the previous species composition. Note that pair-wise differences in the index of monotonicity *m* are proportional to the response dissimilarity *rd* of the same two species. However, response dissimilarity contains more detailed information on differences between the abundance-environment relationships of two species. Accordingly, we use response dissimilarity to calculate response diversity and display response dissimilarities in a two-dimensional ‘response space’ (reflecting pairwise response dissimilarities better than one-dimensional representations) in this study. Nonetheless, conceptual consequences of high or low response diversity are well explainable focusing on *m* denoting rough differences in abundance-environment relationships.

In this study, we exploited performance-environment relationships as species-specific responses to an environmental gradient, using local abundances of plant species as performance proxy (compare to Ruiz-Moreno et al., 2024; Stavert et al., 2017) and land-use intensity as explanatory variable for performance curves as observed at *n* = 150 grassland plots within the Biodiversity Exploratories (Fischer et al., 2010). Response dissimilarity *rd* ∈ [0, 1] quantifies pairwise distances between species-specific abundance-environment relationships described by two empirical checkerboard copulas that are two-dimensional distribution functions that contain full information about associations. The response dissimilarity *rd* matrix (see Fig. 1B) can be used to quantify response diversity of communities analogous to distance-based functional diversity measures (Laliberté & Legendre, 2010; Mouchet et al., 2010; Villéger et al., 2008). Furthermore, the response dissimilarity *rd* matrix can be used to establish a response space where species are positioned according to their (dis-)similarity in performance curves. We decided to use copula-based distances between performance curves as these distribution functions capture full information of species responses to environmental gradients and do not depend on fitting of functions or smoothing parameters (Griessenberger et al., 2022; Junker et al., 2021). Additionally, copulas are not restricted to monotonic function types and thus consider non-monotonic relationships between plant abundance and the environment, which are commonly found in natural systems (Hanusch et al., 2023). For instance, plant species may have their optimum at intermediate land-use intensities (Koch et al., 2016), resulting in non-monotonic performance curves that must be considered in quantifying response diversity and the evaluation of ecosystem stability. Ecosystem stability is a multidimensional property of ecosystems and encompasses several aspects including (among others) a community’s invariance of species diversity and composition or of function and performance (Donohue et al., 2016; Kéfi et al., 2019). Response diversity within functional groups (i.e. functional redundancy of a group of species) is usually associated with positive effects on stability as the functions provided by species suffering from changes will be replaced by species not affected or even facilitated by the same changes (Chillo et al., 2011; Stavert et al., 2017). Here we argue, however, that response diversity in the context of global change with continuous unidirectional shifts in environmental variables may increase functional stability but decrease stability in terms of species diversity and composition. Our predictions as summarized in Fig. 1C stem from the following considerations: In a community consisting of species with divergent responses to environmental change, those species that negatively respond to this change may become weak competitors and eventually will be completely outcompeted by other species better adapted to new conditions (Fig. 1Ci). In contrast, species that positively respond to the same environmental change will become dominant. In this scenario, the species richness will decrease (unstable species composition) and species composition will change in a way that the new communities will be composed of the winners of the change. Thus, the species of the changed community will consist of a set of species nested within the species in the previous community. In contrast, the functionality of the community may remain unaffected (functional stability) if the remaining species compensate for the functions not provided anymore by the lost species (Fig. 1Ci). For instance, ecosystem productivity, either measured directly as biomass production or estimated from spectral reflectance of vegetation (Mazzochini et al., 2024), may be unaffected by species decline if persistent species are functionally redundant to the lost species. Communities consisting of species that collectively respond negatively to the change (low response diversity) may decline in species numbers and also in functionality (Fig. 1Cii). If lost species are replaced by immigrating species better adapted to new conditions, both species number and functionality may be recovered. In that case, the temporal beta-diversity would feature a pronounced turnover component. Finally, in communities consisting of species that collectively respond positively to the change (low response diversity), both species number and composition may be stable and functionality may even increase if species perform better under the new environmental conditions (Fig. 1Ciii).

These scenarios (Fig. 1C) led us to specific hypotheses on the relationship between response diversity, stability, and beta-diversity components that we tested in an experimental common garden setting. The common garden consisted of grass sods from three German regions where plant species diversity and composition has been recorded since 2006 within the framework of the Biodiversity Exploratories (Fischer et al., 2010). Grass sods originated from areas with variable land-use intensity (measured as land-use index LUI, Blüthgen et al., 2012), ranging from protected areas with minimal usage to intensely managed grasslands. Each grass sod was divided in four parts (in the following referred to as subplots) that were subjected to one of four experimental land-use treatments that differ in mowing frequency and amount of fertilizer for three growing seasons. Our hypotheses are:

1) Species respond differently to changes in land-use intensity, and communities in common garden subplots differ in response diversity, resulting from subplot-specific species compositions.
2) Land-use intensity affects response diversity both in the field (long-term) as well as in the common garden (short-term responses).
3) Communities with high response diversity respond to changes in land-use intensity with a decline in species diversity (less stable in diversity).
4) In contrast, communities with high response diversity remain more stable in ecosystem function, e.g. traits indicating productivity.
5) The direction of land-use change (intensification or extensification) affects species composition favoring winners or losers of land-use change and this predicts the direction of shifts within a response space.
6) Temporal beta-diversity is dominated by nested components in response diverse communities, but by turn-over components in communities featuring a low response diversity.

## Methods

### Common garden and experimental procedure

The study was conducted as part of the Biodiversity Exploratories, a large-scale research platform for functional biodiversity research. It comprises three long-term research sites (‘Exploratories’) in Germany at Schwäbische Alb, Hainich, and Schorfheide-Chorin, where land-use and biodiversity are continuously recorded on *n* = 50 grassland plots per region since 2006 (more details about the design can be found in Fischer et al., 2010). Per Exploratory, we selected *n* = 13 plots that cover a large range of land-use intensity (measured as land-use index LUI, see Blüthgen et al., 2012) and collected grass sods with a size of 1m^2^ and 10cm depth in April and May 2020. Each sod was split into four parts of 50×50cm (in the following referred to as subplot) and the subplots were placed in a common garden at the Botanical Garden of the University of Marburg, Germany, to track effects of experimental land-use under reduced environmental heterogeneity. Each of the four subplots per original plot was then assigned to one of four experimental land-use treatments: ‘00’ (= mowing once per year), ‘0M’ (= mowing twice per year), ‘F0’ (= mowing once per year and fertilizer addition) and ‘FM’ (= mowing twice per year and fertilizing). Land-use treatments started in mid-July 2020. Fertilization (99 kg N ha^-1^ year^-1^) and the first mowing of the respective subplots took place in early summer each year, the second mowing in autumn.

The land-use index LUI (Blüthgen et al., 2012), quantifying variation in land-use intensity, was used to account for the land-use history of the plots. The land-use-intensity index (LUI) was calculated as global mean of grassland management for all 3 regions for the years 2006 to 2019 according to Blüthgen et al. (2012), based on information from the land owners on mowing, grazing and fertilization (Vogt et al., 2019) using the LUI calculation tool (Ostrowski et al., 2020) implemented in BExIS (http://doi.org/10.17616/R32P9Q). Grassland plots in the Exploratories had a mean LUI of 1.79 (range: 0.58 – 3.25). Treatments of common garden plots (numbers of cuts [cuts / year] and amount of fertilizer [kg N / ha / year]) were determined based on the 90% quantiles of land-use management data of all plots in the Biodiversity Exploratories between 2006 and 2016 (latest data available at the time of planning) and varied between 0.99 and 2.30 (00 = 0.99, 0M = 1.40, F0 = 2.07, FM = 2.30). Experimental land-use change was quantified by subtracting the mean LUI of plots of origin (Exploratories) from the LUI of common garden subplots, i.e. positive values indicate an increase in land-use intensity, while negative values indicate a decrease in land-use intensity. The taxonomic composition of plant communities in the common garden was recorded in July 2020 (before treatments) and 2022 (after treatments) when all vascular plant species per subplot were identified and the cover of each species (as a proxy for abundance) was estimated with a resolution of 1%. Changes in community composition were calculated as the Bray-Curtis dissimilarity between plot-wise species abundances from 2020 and 2022 using function *vegdist* (R package *vegan* v2.6.4). Shannon diversity of subplots was calculated for 2020 and 2022 with the respective plant species abundance using function *diversity* (R package *vegan* v2.6.4), and the change in Shannon diversity (Δ Shannon) was determined by subtracting the Shannon diversity of 2020 from the Shannon diversity of 2022.

We assessed plant community features through digital whole-community phenotyping (Zieschank & Junker, 2023). Plant communities of subplots in the common garden were scanned with a customized automated plant phenotyping system (PlantEye F500, Phenospex, Heerlen, The Netherlands) that enables the simultaneous recording of multispectral information and the 3-dimensional structure of the vegetation. Scans are then processed with the built-in software HortControl (Phenospex, Heerlen, The Netherlands) that provides *n* = 13 morphological and physiological parameters for further analyses. These digital community features are representative for the frequency distribution of traits and proved to be an effective method to track responses of plant communities to environmental change (He et al., 2023; Zieschank & Junker, 2023). For the analysis we used the scans taken before the first treatment in late July 2020, and those taken after three seasons of applied land-use treatments in mid-August 2022. Changes in plant community features were determined separately for morphological (*n* = 8) and physiological (*n* = 5) community features from digital whole-community phenotyping (see Zieschank & Junker 2023). We calculated Euclidean distances between trait measurements from 2020 and 2022 per subplot using function *vegdist* (R package *vegan* v2.6.4).

### Response dissimilarity *rd* and response diversity *RD*

Performance curves of plant species as basis for the calculation of response dissimilarity *rd* were assessed from local abundances of plant species as performance proxy at *n* = 150 grassland plots within the Biodiversity Exploratories. Only species were considered that occurred on *n* ≥ 25 plots (covering *n* = 96 out of *n* = 149 plant species found in the common garden). Vegetation analysis on Exploratory plots was performed by the core project Botany of the Biodiversity Exploratories. Plant species were identified on a fixed area of 4×4m in May/June from 2008 to 2019 and the abundance of each plant species was estimated (cover in %, REF). The dataset is publicly available on BExIS (https://www.bexis.uni-jena.de/, ID #31389, Version 7). Exploratory plots were ordered by mean land-use intensity index (LUI; values from 2006 to 2019) and we used the R package runner (Kałędkowski, 2023) to calculate a rolling mean of abundances over *k* = 10 consecutive plots for each plant species. Rolling means reflect distribution of taxa across the LUI gradient more realistically as abundance in plots of rather small size (16 m^2^). For each pair of taxa, we calculated the response dissimilarity *rd* to LUI using the R package *qad* (Griessenberger et al., 2021; Griessenberger et al., 2022). For both plant species A and B of a given pair we first computed the mass distribution of the empirical checkerboard copula (function *ECBC*) with rolling mean LUI as explanatory variable and rolling mean abundance of the plant species under consideration as dependent variable resulting in two empirical copulas *A* and *B* with the same resolution (resolution = 12, facilitated by the rolling mean approach). *D1* distance (function *ECBC*) between the copulas *A* and *B* represents a precise estimate of the response dissimilarity *rd* of two taxa to the same environmental gradient. *D1* was appropriately normalized with the constant 2 to yield response dissimilarity *rd* ∈ [0,1]. D1 distances of all species pairs where then compiled into a dissimilarity matrix that was used to position plant species in a ‘response space’ and to calculate response diversity *RD*. To test whether the choice of moving frame size *k* has a significant effect on the results, we performed a Mantel test between distance matrices based on different moving frame sizes *k*. Results showed that distances are highly correlated (correlation between *k* = 5 and *k* = 10: Mantel statistic r: 0.6467, *p* < 0.001 / *k* = 10 and *k* = 25: Mantel statistic r: 0.7079, *p* < 0.001), suggesting no strong effects of choices of *k*.

We used non-metric multidimensional scaling (NMDS) based on the response dissimilarity *rd* matrix using the function *metaMDS* (R package *vegan* v2.6.4) to expand a two-dimensional response space, where the distance between species is proportional to the difference in their responses to land-use intensity. Plant species responses to land-use where classified in either monotonic positive, monotonic negative, or non-monotonic, using the index of monotonicity *m* as described in Hanusch *et al*. (2023), which quantifies the deviation of a given performance curve from ideal monotonicity and returning values in the interval [-1,1]. To determine the direction of association, we used Spearmańs rank correlation to assign *performance curves* to negative monotonic associations (*m* < 0; significant according to Spearmańs rank correlation, ρ < 0), positive monotonic associations (*m* > 0; significant according to Spearmańs rank correlation, ρ > 0), or non-monotonic associations (*m* ∼ 0, not significant according to Spearmańs rank correlation).

Response diversity *RD* was calculated for all subplots of the common garden. Response diversity *RD* was estimated as functional dispersion (FDis) based on the subplot specific abundances of plant species in 2020 or 2022 and the response dissimilarity *rd* matrix. Functional dispersion (FDis) represents the multivariate dispersion weighted by species abundance and is insensitive to species richness and not strongly influenced by outliers (Anderson, 2006; Laliberté & Legendre, 2010). FDis was calculated using the function *dbFD* (R package FD, v1.0.12.1, Laliberté & Legendre, 2010). Changes in *RD* (Δ Response diversity) due to experimental land-use treatments were obtained from subtracting *RD*_2020_ from *RD*_2022_.

### Temporal changes in species composition

To track the magnitude and direction of shifts of plant species from 2020 to 2022 within the response space, we applied vector analysis following the approach described in Junker and Larue-Kontić (2018). Each subplot occupied a fraction of the subplot defined by the position of the plant species present in a given subplot and an abundance-weighted centroid defining the mean position of the species within the response space. The vector between the centroid of a given subplot in 2020 and the centroid of the same subplot in 2022 defined the shift of the plant community. Thus, each vector **u_s_** has the community-weighted mean (cwm) position of the plant species of subplot *s* in 2020 as starting point and the cwm position of the plant species of subplot *s* in 2022 as endpoint. Length *l*(**u_s_**) of vector **u_s_** was calculated by the Euclidean distance between the start- and endpoint of **u_s_**. To compare the direction of the vector among the subplots, we calculated the angle between **u_s_** and a reference vector **u_r_** defined as the vector starting at the origin of the response space and ending at the plant species that most strongly positively responded to land-use changes, *Plantago lanceolata* (highest index of monotonicity *m* = 0.698). The deviation of the unit directions of vectors **u_r_** and **u_s_** in terms of the angle *φ_u_r__,_u_s__* [°] indicates the direction of the shift of a community between 2020 and 2022. *φ_u_r__,_u_s__* = 0° indicates a shift towards plant species with positive responses to land-use (same direction as *P. lanceolata*), *φ_u_r__,_u_s__* = 180° indicates a shift towards plant species with negative responses to land-use.

Additionally to vector analysis, we calculated temporal β-diversity of common garden subplots between 2020 and 2022 and separated its turnover (i.e. replacement of species) and nestedness (species losses) component to test the predictions on the relationship between β-diversity and response diversity. Temporal β-diversity was calculated using Sørensen’s dissimilarity (presence-absence data) of plant species (Baselga, 2010). We partitioned β diversity (β_SOR_) into turnover (β_SIM_) and nestedness (β_SNE_) components using function *beta.temp* in R package *betapart* (v1.6, Baselga et al., 2023). Temporal β diversity was also calculated taking abundance into account, using function *beta.pair.abund* of the same package, measured as Bray-Curtis dissimilarity of subplots between 2020 and 2022. Additionally to the total abundance-based dissimilarity (β_BRAY_), the function also returns values for the balanced variation in abundance (= substitution pattern like species turnover, β_BRAY.BAL_) and the values for the abundance-gradient component of Bray-Curtis dissimilarity (= subset pattern like nestedness, β_BRAY.GRA_) (Baselga, 2013).

To test for relationships between response diversity and other variables characterizing the plant communities in subplots, we fitted linear mixed-effects models using the function *lmer* (R package lme4 v 1.1.31, Bates et al., 2015) and used plot origin (plots in the Exploratories) as random factor to correct for the fact that each original plot was split into 4 subplots. We performed subsequent ANOVA to test for significance using function *anova* (R base package *stats* 4.2.061). All statistical analyses were performed using R: A language and environment for statistical computing (R version 4.2.0, R Core Team, 2022).

## Results

Abundance-environment relationships, i.e. performance curves of plant species present in the common garden were established based on their abundances in *n* = 150 grassland plots of the Biodiversity Exploratories that differed in land-use intensity between 2008 and 2018 (Fig. 2A-C). The dissimilarity of performance curves of two plant species reflects the species’ response dissimilarity *rd*. Based on the response dissimilarity matrix, we conducted a non-metric multidimensional scaling (NMDS) visualizing the response space of *n* = 96 grassland plant species (Fig. 2D). Species’ distribution along the first NMDS axis corresponded to the direction of their association to land-use intensity, i.e. whether they are winners or losers of land use intensification or whether they respond in a non-monotonic way.

**Fig. 2.**
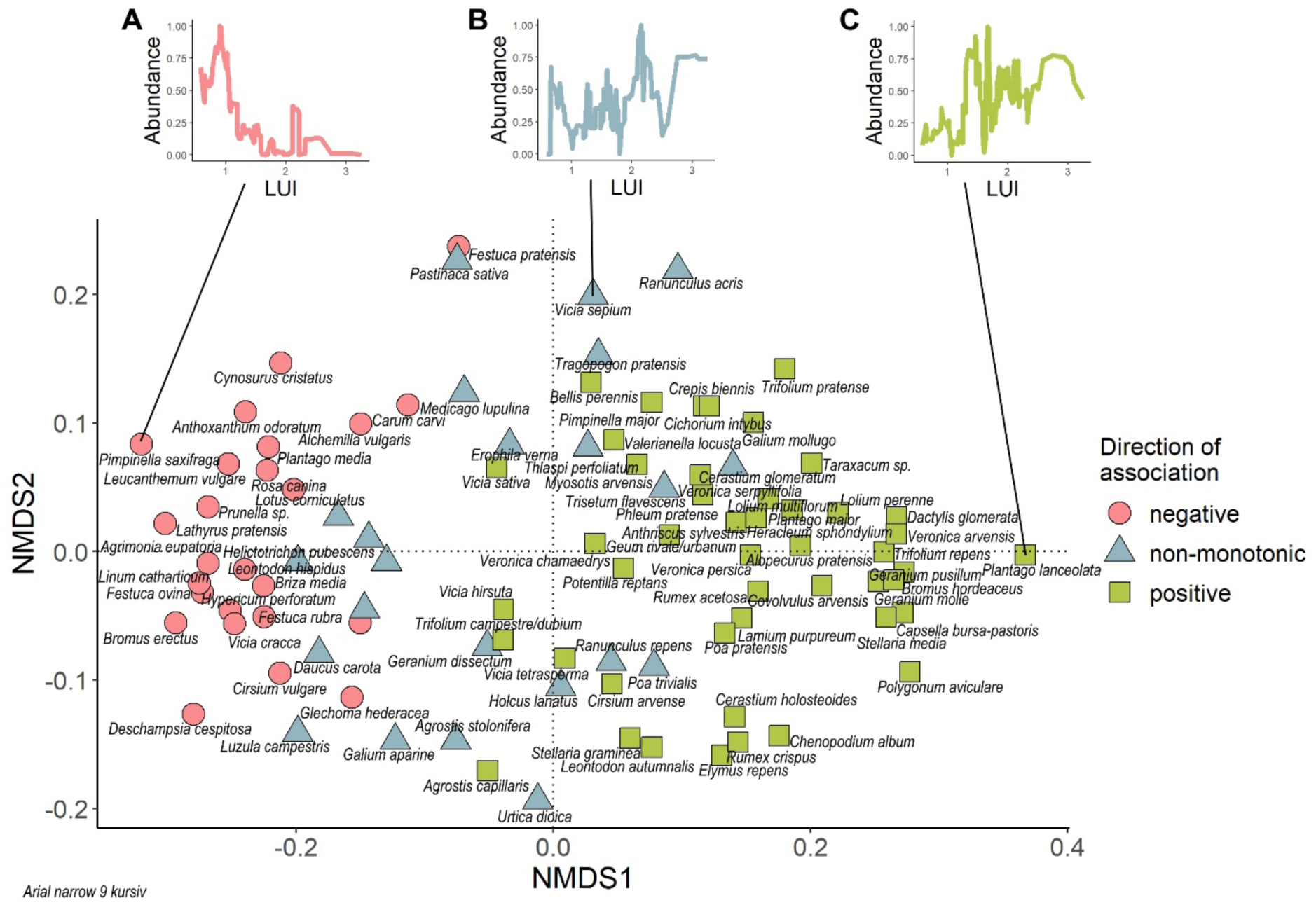
Response-space of grassland plant species to land-use changes. **A-C)** Abundance-environment relationships of three plant species *Pimpinella saxifraga* (A, red, negative response to land use intensification), *Vicia sepium* (B, yellow, non-monotonic response to land use intensification), and *Plantago lanceolata* (C, green, positive response to land use intensification). **D)** Non-metric multidimensional scaling (NMDS) based on response dissimilarities of species pairs separates winners (green circles, positive indices of monotonicity *m*) from losers (red circles, negative indices of monotonicity *m*) of land-use intensification as well as those species that show non-monotonic responses to land-use intensity (yellow circles, indices of monotonicity *m* ranging around zero). Note that pair-wise differences in the index of monotonicity *m* are proportional to the response dissimilarity *rd* of the same two species (Mantel statistic r: 0.8067, p < 0.001).

In a common garden, we subjected plant communities that originated from different plots within the Biodiversity Exploratories (sod transplantation) to four land use treatments representing a range of land use intensities (LUI). The response diversity *RD* of plant communities at the beginning of the experiment showed a pronounced variation and was negatively related to the historical land-use intensity on the plots they originated from (linear mixed model with plot origin as random factor and subsequent ANOVA: *F*_1,37_ = 5.68, *p* = 0.022; Fig. 3A). In contrast, initial Shannon diversity of plant communities showed no significant relationship to land-use history of the plots they originated from (*F*_1,37_ = 3.81, *p* = 0.0585; Supplementary Fig. 1A). After three seasons of experimental land-use treatments, response diversity RD of plant communities in the common garden decreased with positive changes in land-use intensity and increased in subplots with negative changes in land-use intensity (linear mixed model with plot origin as random factor and subsequent ANOVA: *F*_1,184_ = 4.43, *p* = 0.037; Fig. 3B), which is in line with the negative relationship between response diversity RD and the land-use history in the field (Fig. 3A). Shannon diversity declined drastically on subplots where land-use was intensified (linear mixed model with plot origin as random factor and subsequent ANOVA: *F*_1,154_ = 17.74, *p* < 0.001; Supplementary Fig. 1B), but only barely increased with decreasing land-use intensity. In line with our predictions (Fig. 1C), Shannon diversity strongly decreased in communities with a high initial response diversity RD and increased in communities with a low initial response diversity RD within the course of our experiment (linear mixed model with plot origin as random factor and subsequent ANOVA: *F*_1,80_ = 42.80, *p* < 0.001; Fig. 3C). Likewise, in the first year of the experiment, community composition changed more strongly in communities with high initial response diversities compared to those with low initial response diversities (linear mixed model with plot origin as random factor and subsequent ANOVA: *F*_1,110_ = 4.18, *p* = 0.043). The following years, this trend was still visible, although not significant (linear mixed model with plot origin as random factor and subsequent ANOVA: 2021: *F*_1,69_ = 1.91, *p* = 0.171; 2022: *F*_1,94_ = 2.33, *p* = 0.130). In contrast to the lower stability in species diversity in plant communities with high response diversities, the functionality remained more stable (weaker changes in physiological multivariate community features) in communities with high initial response diversity RD (*F*_1,151_ = 4.91, *p* = 0.028; Fig. 3D). We found no significant relationship between the strength of changes of morphological community features and the response diversity RD of plant communities (*F*_1,62_ = 0.07, *p* = 0.788).

**Fig. 3.**
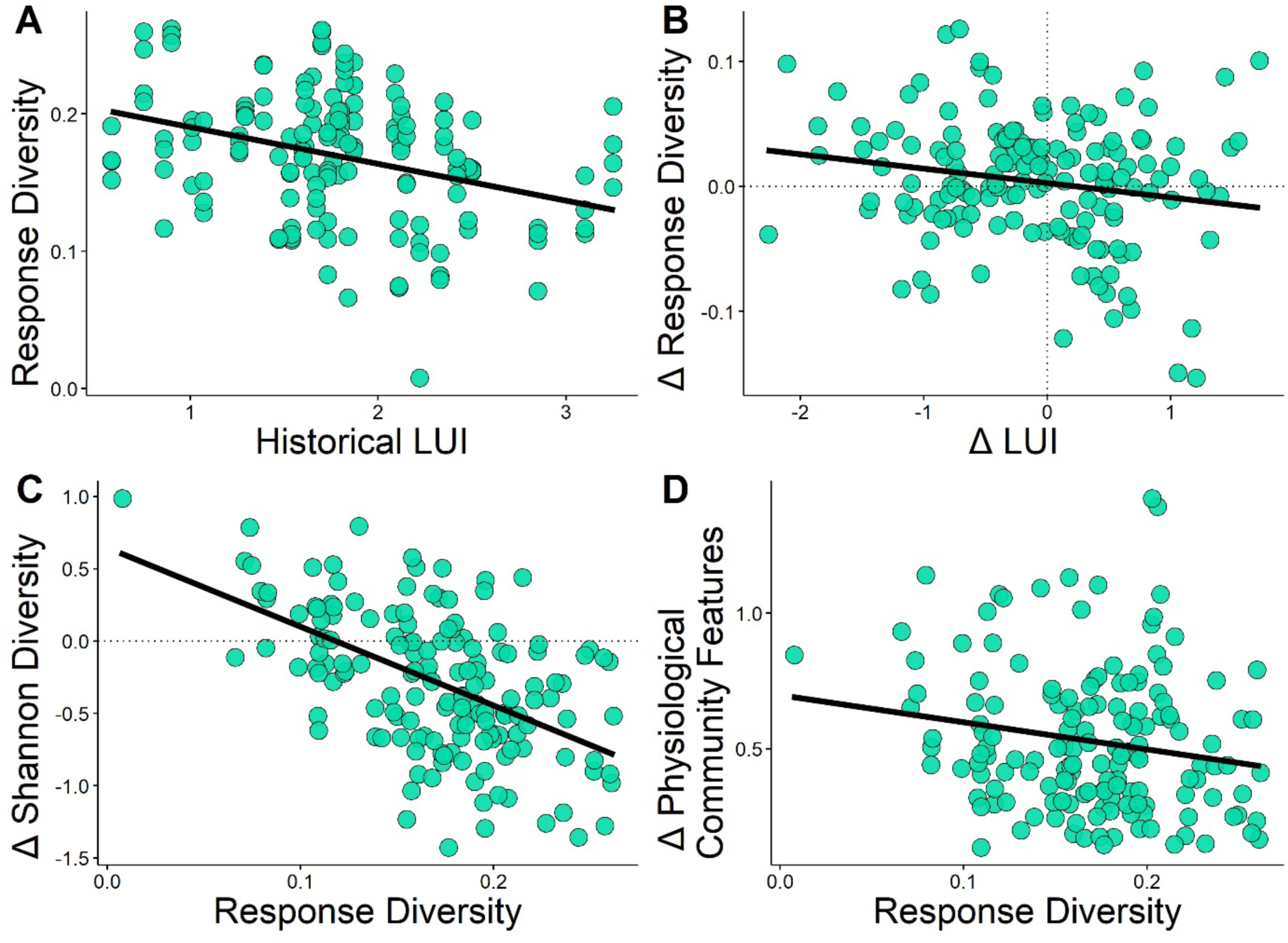
Community responses to land-use changes in the field and the experimental plots as a function of response diversity RD. **A)** Negative relationship between the historical land-use intensity (LUI) and the response diversity RD of communities in the field. **B)** Negative relationship between experimental changes in land-use intensity (Δ LUI) and changes in the response diversity RD (Δ response diversity) in the course of the experiment. **C)** Plant communities with high initial response diversities decreased in Shannon diversity in the course of the experiment. **D)** Plant communities with high initial response diversities had little changes in physiological community features in the course of the experiment and thus remained functionally more stable than plant communities with low initial response diversity RD.

To track the strength and direction of shifts of plant communities within the response space (Fig. 2), and course of our experiment, we quantified the lengths *l*(**u_s_**) of vectors **u_s_** with the community-weighted mean (cwm) position of the plant species of subplot *s* in 2020 as starting point and the cwm position of the plant species of subplot *s* in 2022 as endpoint. Vector length *l*(**u_s_**) positively correlated with the change in community composition quantified as Bray-Curtis distance based on plant species composition in each subplot recorded in 2020 and 2022 (linear mixed model with plot origin as random factor and subsequent ANOVA: *F*_1,138_ = 45.25, *p* < 0.001). Vector length *l*(**u_s_**) was positively correlated to changes in response diversity RD within in the course of the experiment (combined linear mixed model with plot origin as random factor and subsequent ANOVA: *F*_1,151_ = 7.09, *p* = 0.009) but negatively with changes in Shannon diversity (combined linear mixed model with plot origin as random factor and subsequent ANOVA: *F*_1,143_ = 4.92, *p* = 0.028, Fig. 4A). Additionally, we quantified the angles *φ_u_r__,_u_s__* [°] between the vectors **u_s_** and a reference vector **u_r_** defined as the vector starting at the origin of the response space and ending at the plant species that most strongly positively responded to land-use changes, *Plantago lanceolata* (highest index of monotonicity *m* = 0.698). *φ_u_r__,_u_s__* = 0° indicates a shift towards plant species with positive responses to land-use (same direction as *P. lanceolata*), *φ_u_r__,_u_s__* = 180° indicates a shift towards plant species with negative responses to land-use. Vector angle *φ_u_r__,_u_s__* was negatively correlated to the change in land-use intensity (linear mixed model with plot origin as random factor and subsequent ANOVA: *F*_1,144_ = 4.48, *p* = 0.036, Fig. 4B), showing that plant communities subjected to increased land-use intensity shifted towards the winner-species within the response space (low *φ_u_r__,_u_s__*, Fig. 4C), whereas plant communities subjected to decreased land-use intensity shifted towards losers (high *φ_u_r__,_u_s__*, Fig. 4D).

**Fig.4.**
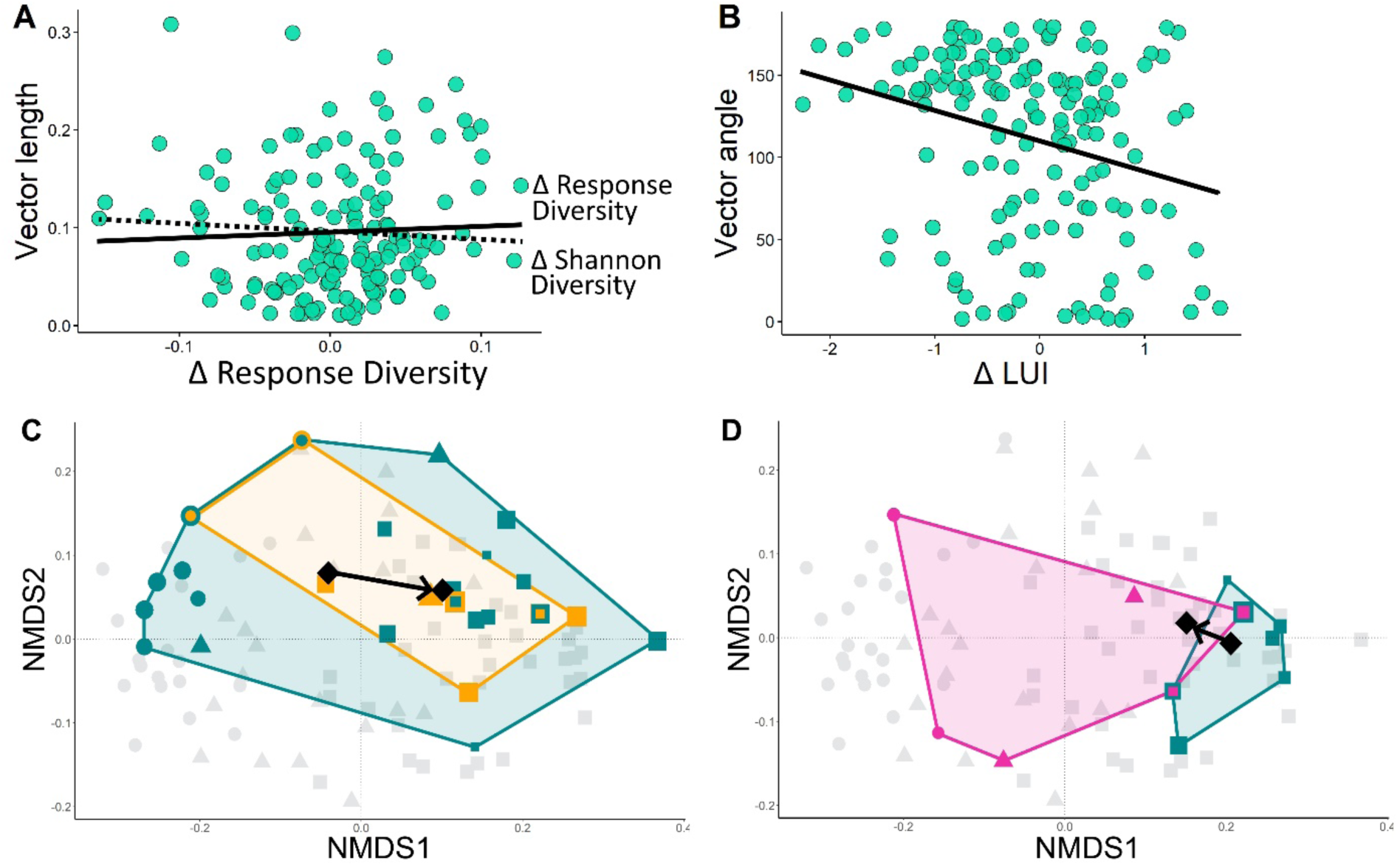
Strength and direction of community shifts within the response space after experimental land use. **A)** Negative relationship between experimental changes in land use intensity and the angle *φ_u_r__,_u_s__* of vectors of community shifts within the response space to the reference vector pointing towards the most positive response to land use intensification. Low *φ_u_r__,_u_s__* denote shifts towards winner species; high *φ_u_r__,_u_s__* denote shifts towards loser species. **B)** Relationship between the changes in response diversity RD or Shannon diversity in response to experimental land-use change and the vector length *l*(**u_s_**). **C)** Example for plant community changes in response to experimental land-use intensification. Each symbol represents a plant species. Blue species (and blue convex hull) were present in 2020, symbol size is proportional to relative abundance of species. Yellow species (and yellow convex hull) were present in 2022 after experimental land-use change. Blue species with yellow frames were present throughout the experiment. Grey plants species are not present in the focal plant community but serve as a reference for the response space. Symbol shapes indicate plant species responses to land-use change with circles denoting negative, squares denoting positive and triangles denoting non-monotonic responses. Black symbols denote community weighted mean positions, i.e. centroids of plant species’ positions within the response space, at the beginning and the end of the experiment, the vector indicates the length and direction of shift. **D)** Same as in C but an example for plant community changes in response to experimental reduction in land-use. Blue species (and blue convex hull) were present in 2020, pink species (and pink convex hull) were present in 2022 after experimental land-use change.

To gain insight into the processes of community shifts in response to experimental land-use changes as a function of initial response diversity RD, we decomposed the temporal β diversity of each subplot into a turnover (= replacement of species) and nestedness (= species losses) component. β diversity was calculated with the Sørensen index considering the presence/absence of species and revealed that the nestedness component increased with increasing initial response diversity RD of plant communities (Fig. 5, linear mixed model with plot origin as random factor and subsequent ANOVA: *F*_1,73_ = 16.18, *p* < 0.001), indicating that experimental land-use changes led to reduced numbers of species without replacement in communities with high response diversity RD. As the nestedness + turnover = overall β diversity, the turnover component was negatively correlated to initial response diversity RD (linear mixed model with plot origin as random factor and subsequent ANOVA: *F*_1,81_ = 9.74, *p* = 0.002), indicating a replacement of species (immigration) in subplots with low response diversity RD. Overall temporal β diversity of plant communities did not correlate with response diversity RD (linear mixed model with plot origin as random factor and subsequent ANOVA: *F*_1,89_ = 1.18, *p* = 0.280). Temporal β diversity based on Bray-Curtis distances, i.e. considering also species abundances, was unrelated to initial response diversity RD (linear mixed model with plot origin as random factor and subsequent ANOVA: *F*_1,94_ = 2.21, *p* = 0.140). The same is true for nestedness and turnover components based on Bray-Curtis distances (linear mixed model with plot origin as random factor and subsequent ANOVA: turnover: *F*_1,85_ = 2.20, *p* = 0.142; nestedness: *F*_1,59_ = 0.01, *p* = 0.951).

**Fig. 5.**
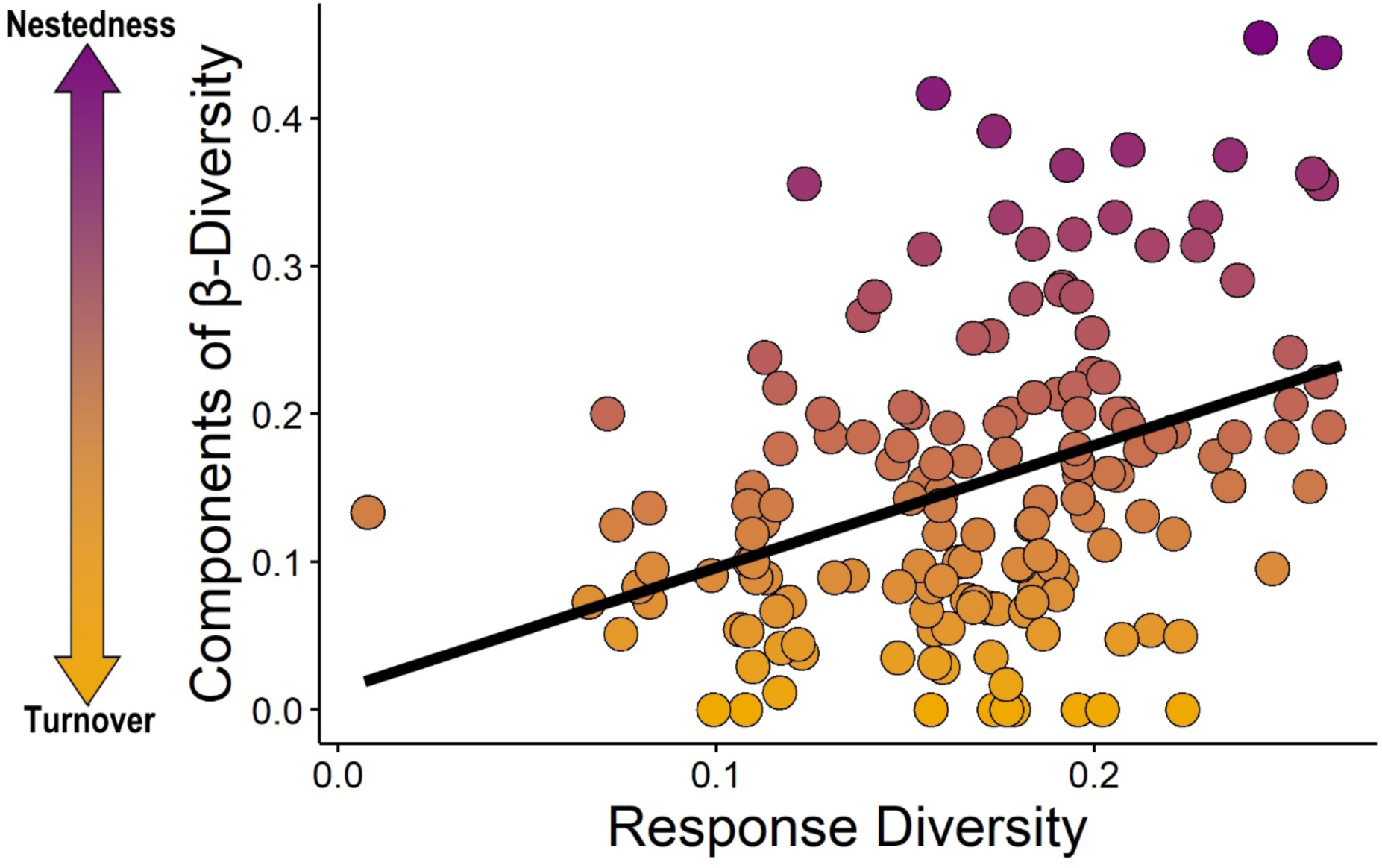
Temporal β diversity of community composition in response to experimental land use changes as a function of initial response diversity RD. Temporal β diversity of each subplot was decomposed into a turnover (= replacement of species) and nestedness (= species losses) component. β diversity was calculated with the Sørensen index considering the presence/absence of species revealed that the nestedness component increased with increasing initial response diversity RD, whereas the turnover component decreased in with initial response diversity RD. These findings together with those presented in Fig. 4 indicate that plant communities featuring a high response diversity RD often loose species (likely those with positive responses to land-use changes), whereas communities with a low response diversity RD often change community composition by favoring immigrating species that are better adapted to new conditions.

## Discussion

Our common garden experiment revealed causes and consequences of response diversity among grassland species in the context of anthropogenic land-use changes. Our six hypotheses on the interplay between response diversity, land-use change and plant community responses were largely supported by our results. (H1) Abundance-environment relationships of common grassland species varied substantially, ranging from negative over non-monotonic to positive responses to land-use intensification, which translated into species-specific positions in a response space. Plot-specific compositions of species that featured either similar or dissimilar responses resulted in low or high response diversity, respectively. (H2) Our field and experimental common garden data indicate a negative effect of land-use intensification on response diversity, which means that high land-use intensity favors plant species with homogenous responses to land-use changes. (H3 and H4) We found that plant communities with high response diversities are less stable in terms of species diversity, but more stable in terms of functionality than plant communities with a low response diversity. (H5) Land-use changes additionally affected plant species composition in a way that land-use intensification favored plant species that positively respond to land-use intensification, whereas land-use extensification favored plant species that negatively respond to land-use intensification. (H6) Finally, land-use changes caused alterations in species compositions in dependence of initial response diversity: communities with an initially high response diversity lost species that were not replaced by immigrating species (see H3), resulting in a dominant nested component of temporal beta-diversity. In contrast, communities with an initially low response diversity also lost plant species, but those were then replaced by immigrating species, which becomes apparent in a strong turnover component of temporal beta-diversity. Yachi and Loreau (1999) identified the form and degree of asynchronicity of individual species’ responses to environmental fluctuations as determinants for the insurance effects of species-rich communities. In their model, species-rich communities featured less variable and on average higher functionality than species poor communities in fluctuating environments. These effects were attributed to asynchronous species-specific variation in the provisioning of certain functions, but not the complete loss of species and their contributions to functionality. In times of global change, environmental conditions often do not only fluctuate, but constantly shift in one direction with consequences not only for quantitative measures of species’ functions, but also for their persistence (presence / absence) at local and regional scales. For instance, land-use intensification leads to changes in productivity (Allan et al., 2015; Schils et al., 2022) and also in species composition and diversity (Gossner et al., 2016; Weiner et al., 2011), which challenges communities and ecosystems in different and potentially more severe ways than temporal fluctuations. Our study confirmed these effects by showing clear community responses to an increased land-use intensity that affect the temporal stability of communities both in the field as well as in our experimental grassland plots.

While theoretical and conceptual work on the insurance hypothesis and response diversity usually addresses functional stability (Mori et al., 2013; Yachi & Loreau, 1999), empirical studies considered additional measures for stability that reflect the multidimensionality of this ecosystem property (Donohue et al., 2016). These measures for stability included the variance of functional diversity based on response traits (Aquilué et al., 2020; Laliberte et al., 2010), the persistence of species and thus species composition and diversity (Baskett et al., 2014; Chillo et al., 2011; Ives et al., 1999; Moore & Olden, 2017), quantitative measures of ecosystem functions (Cariveau et al., 2013; Moore & Olden, 2017; Sasaki et al., 2019), as well as proxies for ecosystem functions (Mazzochini et al., 2024). Our study revealed that response diversity in the context of global change with continuous unidirectional shifts in environmental variables stabilized ecosystem functions, suggesting that plant species with positive performance-environment relationships compensated for the loss of species and their functionality, which is fully in line with the insurance hypothesis of biodiversity (Yachi & Loreau, 1999). In contrast, high response diversity destabilized plant communities in terms of species composition and diversity as the experimental unidirectional shifts in environmental variables in our common garden experiment favored the winners of land-use change, while losers were replaced. Our data reflect real-world results, separating plant species into those that respond negatively, positively, or do not show a monotonic response to a land-use gradient (Busch et al., 2019), which leads to a decrease in diversity and a homogenization of plant communities with increasing land-use (Gilhaus et al., 2017; Gossner et al., 2016; Kachler et al., 2023; Socher et al., 2013). The response space as introduced in in this study helps to identify species that are particularly threatened by global change components and also groups of species with similar responses. Both information may help to evaluate the response diversity and stability of communities and ecosystems. The results presented here may be more accentuated than those of field studies as common gardens reduce environmental heterogeneity and thus help to study effects of the experimental treatments in isolation. Future studies should evaluate the results presented here under more natural conditions with multiple global change components acting simultaneously on vegetation stability, which may alter effects on grasslands (Bütof et al., 2012; De Chazal & Rounsevell, 2009; Pompe et al., 2008).

Grasslands in Central Europe are part of a cultural landscape created by human intervention (Leuschner & Ellenberg, 2017; Poschlod, 2017) and require human management for their conservation (Bucharova et al., 2020; Squires et al., 2018). Therefore, even the plots with the lowest land-use intensity in our study are not pristine ecosystems and the long-lasting human interference has already had a lasting impact on the diversity and other biological characteristics of these areas (Finderup Nielsen et al., 2021; Scherreiks et al., 2022; Wesche et al., 2012). Thus, neither the field data nor the experimental common garden plots cover the full land-use gradient and lack the lower end of the continuum, which could explain the relatively high number of species that show a positive response to land-use. Accordingly, we may assume a bell-shaped relationship between land-use intensity and response diversity with the extremes (very low and very high land-use), supporting only low response diversities. Therefore, the negative relationship between land-use intensity and response diversity found in our study may only represent the left part of the this assumed curve.

Our results, along with variable reported net effects of response diversity on ecosystem stability ranging from positive (Aquilué et al., 2020; Sasaki et al., 2019) to no effects (Cariveau et al., 2013; Elsalahy et al., 2020), indicate that the stabilizing effect of response diversity may be system-specific and clearly depends on how stability is defined and measured (compare to Fig. 1). Furthermore, and maybe most important, the nature of environmental change may strongly affect the relationship between response diversity and ecosystem stability. For instance, temporal fluctuations where environmental conditions alternately deviate positively and negatively from a mean condition (as assumed in the insurance hypothesis of biodiversity, see Yachi & Loreau, 1999) exert other stresses to ecosystems than pulses or unidirectionally shifts in environmental parameters as currently observed in many global change components (Donohue et al., 2016; Kéfi et al., 2019). Additionally, as our data demonstrate, the magnitude of change clearly affects stability and the buffering effect of response diversity (Donohue et al., 2016). Finally, most studies, including ours, focus on a single changing environmental factor or stress (Donohue et al., 2016; Kéfi et al., 2019; Ross et al., 2023). Most ecosystems, however, are challenged by changes in multiple factors or multifactorial stress combinations that may act synchronously or sequentially (Zandalinas & Mittler, 2022). It is questionable whether previously stressed communities with consequently reduced response diversities are still sufficiently resistant or resilient to cope with future stresses. In this context, future studies may specifically test whether species tolerant against one stressor are also tolerant against another stressor, or whether species feature tolerance against a specific set of stressors while they cannot tolerate others. If species can be divided into those with a universal tolerance and those with rather low thresholds of stress tolerance, communities may quickly loose considerable components of their biodiversity.

We conclude that response diversity is an important determinant for ecosystem stability and thus an important target for conservation efforts. Performance-environment relationships turned out to be a powerful tool for assessing pairwise response dissimilarities and response diversity that do not depend on proxies for species responses to environmental changes (compare to Ross et al., 2023; Ross & Sasaki, 2024). However, the strong context dependency of response diversity – stability relationships require system-specific definitions of responses as well as of stability in order to adequately measure and take full advantage of this dimension of diversity. Further insights into the role of multifactorial stress combinations as well as the nature and magnitude of change in affecting response diversity and ecosystem stability are clearly required to develop a robust conceptual framework, and to design conservation strategies that facilitate resilience and resistance and thus maintain ecosystem functioning.

## Data availability

This work was performed within the scope of the Biodiversity Exploratories program (DFG Priority Program 1374). Long-term field data was recorded by the Botany Core project of the Biodiversity Exploratories program (DFG Priority Program 1374) and is publicly available in the Biodiversity Exploratories Information System (http://doi.org/10.17616/R32P9Q). The dataset is listed in the references section (Hinderling & Keller, 2023). Datasets on common garden plant communities have been generated within the project EXClAvE and are also publicly available in the Biodiversity Exploratories Information System (Id 31267 & 31269).

## Acknowledgements

We thank the Botanical Garden of the University of Marburg for their support in establishing and maintaining the common garden, David Meyer, Lilian Winzer, Marie Englert and Marlene Hölzer for their help with field work and data collection, and Annette Schriever and Alexander Lach for technical support. We thank the managers of the three Exploratories, Julia Bass, Anna K. Franke, Miriam Teuscher and all former managers for their work in maintaining the plot and project infrastructure; Christiane Fischer and Victoria Grießmeier for giving support through the central office, Andreas Ostrowski for managing the central data base, and Markus Fischer, Eduard Linsenmair, Dominik Hessenmöller, Daniel Prati, Ingo Schöning, François Buscot, Ernst-Detlef Schulze, Wolfgang W. Weisser and the late Elisabeth Kalko for their role in setting up the Biodiversity Exploratories project. We thank the administration of the Hainich national park, the UNESCO Biosphere Reserve Swabian Alb and the UNESCO Biosphere Reserve Schorfheide-Chorin as well as all land owners for the excellent collaboration. Field work permits were issued by the responsible state environmental offices of Baden-Württemberg, Thüringen, and Brandenburg.

## Funding

The work has been funded by the DFG Priority Program 1374 ‘Biodiversity Exploratories’ (JU 2856/4-1).

## Competing interests

The authors declare that the research was conducted in the absence of any commercial or financial relationships that could be construed as a potential conflict of interest.

## Author contributions

RRJ conceived the study, VZ and RRJ designed the study, VZ performed field work, VZ and RRJ performed statistical analysis, VZ and RRJ wrote the manuscript.

## Supplementary Information

**Supplementary Fig. 1.**
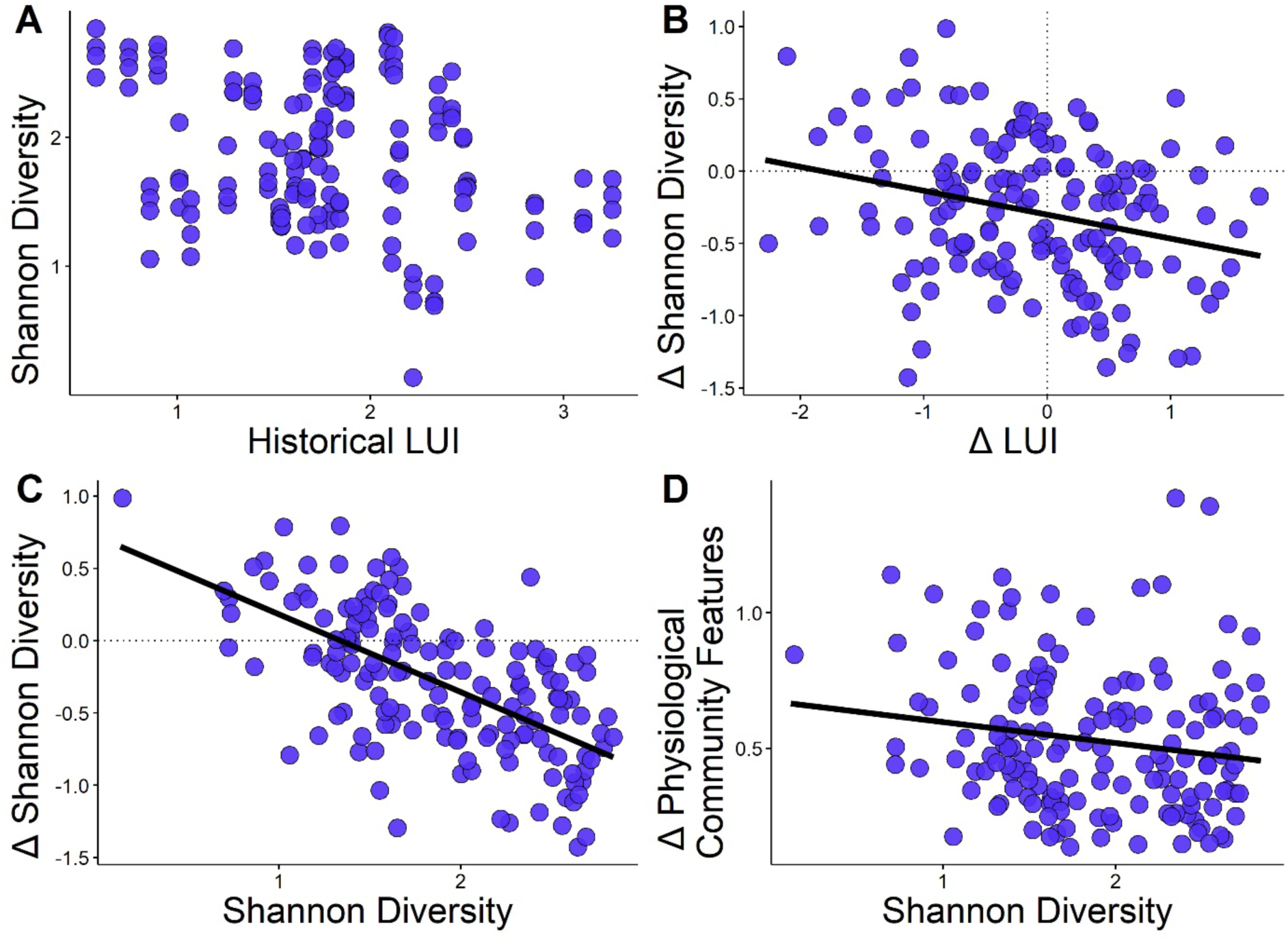
Community responses to land-use changes in the field and the experimental plots as a function of Shannon diversity RD. **A)** Non-significant relationship between the historical land-use intensity (LUI) and Shannon diversity of plant communities in the field. **B)** Negative relationship between experimental changes in land-use intensity (Δ LUI) and changes in Shannon diversity (Δ Shannon diversity) in the course of the experiment. **C)** Plant communities with high initial Shannon diversities decreased in Shannon diversity in the course of the experiment. **D)** Negative relationship between Shannon diversity and changes in physiological community features in the course of the experiment.

## Notes

### Competing Interest Statement

The authors have declared no competing interest.

